# Precision Imaging for Intraindividual Investigation of the Reward Response

**DOI:** 10.1101/2025.09.26.678878

**Authors:** Matthew Mattoni, Shenghan Wang, Cooper J Sharp, Thomas M Olino, David V Smith

**Affiliations:** Temple University, Department of Psychology and Neuroscience

## Abstract

The reliance of fMRI research on between-person comparisons is limited by low test-retest reliability and inability to explain within-person processes. Intraindividual studies are needed to understand how changes in brain functioning relate to changes in behavior. Here, we present open data and analysis of a novel intensively sampled fMRI study, the Night Owls Scan Club. This precision imaging dataset includes 44 sessions acquired across four participants at a twice-weekly rate. In each session, participants completed multiple reward-related tasks, mood and alertness ratings, and a behavioral mood manipulation. We examined how the reward response reflects between-person or within-person variance. Test-retest-reliability of the reward response was very low and not explained by measurement error, suggesting little utility for between-person comparisons. At an intraindividual level, the mood induction showed small increases in the reward anticipation response. Additionally, mood and alertness explained notable intraindividual variance of the reward response, including as much as 31% for one participant. Overall, results suggest that BOLD activation to reward tasks – and likely other fMRI tasks – is more appropriate for within-person study than between-person study, highlighting a need for intensive longitudinal neuroimaging designs.

## Introduction

Neuroimaging research mostly relies on cross-sectional studies and behavioral associations are studied through tests of individual differences. While this between-person focus is critical for goals such as biomarker discovery, statistical power is limited by low test-retest reliability, especially in task-based designs. Moreover, many inferential goals in neuroimaging, such as causes or consequences of changes in brain functioning, reflect within-person processes. As between-person results cannot inform within-person processes, there is a large need for intraindividual fMRI research (Hunter et al., 2024; Mattoni, Fisher, et al., 2025). To address this need, we present an open, intensively sampled dataset and analysis focused on the BOLD reward response. Our primary goals were to (1) assess the test-retest reliability of the reward response, (2) examine state-like factors associated with within-person changes in the reward response, and (3) examine how precision imaging methods impact the measurement of the reward response. To address these aims, we obtained 44 sessions of fMRI data across four individuals, acquired at a roughly biweekly assessment schedule using multiband multi-echo sequencing. In each session, participants completed two runs each of two reward tasks, provided multiple ratings of mood and alertness, and completed a positive mood induction behavioral manipulation in between task runs.

Reward functioning supports fundamental behaviors like decision making (Knutson et al., 2007) and motivation (Bromberg-Martin et al., 2010) and is implicated in numerous clinical disorders (Zald & Treadway, 2017). Subsequently, tasks that target reward processes like the monetary incentive delay (MID) task (Knutson et al., 2000) are some of the most common in fMRI studies. Like in most neuroimaging studies, reward-related studies typically examine individual differences: between-person associations of the BOLD response to reward and targeted behaviors like depression scores. However, the validity of these tests depends on reliability. In fMRI, reliability is frequently considered in terms of test-test-retest reliability, typically measured by intraclass correlation coefficient (ICC; see Noble et al. (2019) for review). However, most evidence suggests that the test-retest reliability of most BOLD task responses, especially the reward response, is low. In a meta-analysis of common fMRI tasks (N > 1000), Elliott et al. (2020) found the mean test-retest reliability to be near .40 (poor) across tasks, including ICCs of .14, .29, .39 (Fliessbach et al., 2010), .28 (Chase et al., 2015), .58 (Holiga et al., 2018), and .59 (Plichta et al., 2012) for reward tasks. More recently, Demidenko et al. (2024) conducted a multiverse analysis of different analytical pipelines for test-retest reliability of the MID task in several large studies and found the ICC to be below .10 for the nucleus accumbens, a common striatal region of interest for reward studies.

Low test-retest reliability suggests that the BOLD reward response may not be appropriate for tests of individual differences (Elliott et al., 2020), evidenced by low statistical power. For instance, with optimistic assumptions of the neuroimaging and behavioral measures each having a reliability of .80, reaching 80% power at an effect size of r = .30 and α = .05 would only require a sample of N = 53. By solely changing the test-retest reliability of the BOLD response to .30, the necessary sample size increases to N=361. Assuming a BOLD response test-retest reliability of .10 found by Demidenko et al. in the nucleus accumbens, the necessary sample size balloons to N=1087. Thus, due to low test-retest reliability alone, reliable tests of individual differences may require thousands of individuals (compounding consequences of small effect sizes; see Marek et al., 2022).

In the context of between-person comparisons, low test-retest reliability is a major obstacle and is generally viewed as a product of measurement error. Studies have examined factors such as retest interval length, analytical decisions, and data quality that may decrease reliability. However, there is minimal evidence of effects of interval length (Elliott et al., 2020; though conclusions are tempered by little data examining sessions days or weeks apart) or analytical decisions (Demidenko et al., 2024). Data quality measures such as signal-to-noise ratios and head motion have also shown little impact on reward response test-retest reliability (Chase et al., 2015).

Moreover, analytical choices that improve test-retest reliability are not inherently more appropriate. For example, two methods with modest improvements to test-retest reliability – larger smoothing kernels and models that contrast reward responses to an “implicit baseline”, rather than a control condition like neutral feedback (Demidenko et al., 2024) – reduce neural and behavioral interpretability, respectively. In sum, with goals of testing individual differences, low test-retest reliability provides a pessimistic view for the utility of the reward response and other task-based responses (Elliott et al., 2020).

However, with an inferential focus on *within-person* variance, low test-retest reliability instead presents an opportunity as a potential signal of true change. For instance, fluctuations in reward response across time may reflect within-person changes in factors affecting how rewarding a stimulus may be, such as mood, interest, or alertness. Importantly, within-person associations reflect distinct processes than between-person associations. While tests of individual differences can inform broad group differences and biological risk markers, within-person associations reflect more mechanistic processes and causal inferences (Hamaker, 2012). For example, while it has been well established (Forbes & Dahl, 2012) that individuals with depression have blunted reward responses (between-person), little is known about what changes in the brain leading to the development or treatment of depression (within-person). Moreover, within-person information has distinct practical implications (Gell et al., 2024; Mattoni, Fisher, et al., 2025). Even if a reliable clinical biomarker was identified, there are substantial barriers to widespread diagnostic use such as scanning time, costs, and relative benefit to other measures. On the other hand, within-person inferences could inform translational research, such as intervention development (e.g., causal study of an intervention altering brain functioning leading to symptom reduction). Moreover, longitudinal designs can improve between-person study by disaggregating within-person variance (Hamaker, 2023; e.g., does a participant have a consistently blunted response to reward, or are they in a particularly unmotivated state?).

Despite the theoretical and practical importance of within-person inferences, between-person research dominates the field. It is thus common to transfer between-person results for within-person inferences (Cragg et al., 2019). However, this assumption of group-to-individual generalizability is unfounded for nearly all human processes (Fisher et al., 2018; Molenaar, 2004), including brain functioning (for review, see Mattoni et al., 2025). Concretely, we cannot assume from findings that individuals with a blunted reward response are more prone to depression (between-person), that an individual’s reward response decreases when they become more depressed (within-person). Relationships at different levels of analysis rely on distinct information (Cattell, 1988) and processes, and effects could be similar or entirely opposite (i.e., “Simpson’s Paradox; Kievit et al., 2013). Thus, within-person conclusions can only be reached through within-person study designs (Mattoni, Fisher, et al., 2025).

As within-person study designs are scarce in fMRI, we have little understanding of intraindividual brain-behavior relationships. Even in studies that collect two or three measurements, within-person inferences are still limited by statistical power (based on number of observations rather than individuals), long intervals between observations, and group-level effects that may not reflect individuals (Hunter et al., 2024). Precision imaging approaches that repeatedly scan individuals are a promising approach to address this gap (Lee et al., 2024). Precision imaging largely grew from demonstrations that longer fMRI time series were needed for reliable individual-level resting state functional connectivity networks (Gordon et al., 2017; Kraus et al., 2023; Laumann et al., 2015). Though large-scale resting state networks show mostly trait-like stability across sessions, appreciable variance is explained by task state and task-evoked signals show much less stability (Gratton et al., 2018). Often as a secondary focus, precision studies have provided key insights into within-person variation and how it differs from between-person variation, even in the same measures (Gratton et al., 2018; Lynch et al., 2024). However, there has so far been little attention to systematically examining brain-behavior relationships over time. In one exception, Flournoy et al. (2024) followed a 30-person sample using an emotional processing fMRI task each month for a year. They found test-retest reliability was very low across the brain – with ICCs approaching 0 – suggesting the task was inappropriate for between-person study. Reliability was not low due to noise, as the task response showed consistent voxel activation across sessions. Instead, within-person fluctuations were explained by intraindividual variation in mood, sleep, and stress. Overall, Flournoy et al. (2024) provided compelling evidence that their emotional processing task was ideal for within-person study, not between-person. There findings are notably consistent with prior evidence that, relative to functional connectivity networks, task-evoked signals are less stable and more reflective of state contexts (Gratton et al., 2018). Thus, task-based activations may be particularly promising for within-person study.

Here, we present the Night Owls Scan Club (NOSC), a reward focused, intraindividual extension of precision imaging study designs (Flournoy et al., 2024; Gordon et al., 2017). Our overall goal was to assess whether the BOLD reward response is trait-like with measurement error or state-like with intraindividual associations. We scanned four individuals twice weekly for up to 12 sessions using multiband multi-echo sequences, with each session consisting of two runs of two different reward tasks and a mood induction experimental manipulation. Trait-like properties were mainly operationalized by test-retest reliability, with secondary exploration of variance across sessions. With an intensive observation schedule, we addressed concern that test-retest reliability is limited by long intervals. We also assessed whether low reliability was explained by measurement noise by using single-trial models to assess the split-half reliability of individual task runs. For intraindividual relationships with behavior, we focused on associations of mood and alertness with the reward response due the relevance of reward functioning to mood (Nusslock & Alloy, 2017), effects of alertness on intraindividual changes in functional connectivity (Laumann et al., 2017; Tagliazucchi & Laufs, 2014), and their intraindividual associations found in Flournoy et al. (2024). Finally, we assessed whether precision imaging methods of multi-echo sequencing and denoising and multivariate reward signatures impacted results. Overall, NOSC presents high-quality precision imaging data with an intensive sampling schedule and experimental manipulations, enabling more detailed study of trait and state properties of the reward response.

## Methods

We collected and publicly released data for the Night Owls Scan Club (NOSC), a precision imaging study of the neural reward response. NOSC includes four individuals scanned numerous times in a short period, producing nearly 200 runs of functional data in total that are accompanied by mood and alertness self-report ratings. Raw BIDS data are available on OpenNeuro (https://doi.org/10.18112/openneuro.ds006707.v1.1.1; accession number ds006707; Mattoni, Wang, et al., 2025), derivatives used for this manuscript are available on Open Science Framework (OSF; https://osf.io/df5t9/; DOI 10.17605/OSF.IO/DF5T9), and code to reproduce all analyses is available at https://github.com/DVS-Lab/night-owls.

### Procedures

We recruited four participants for the study who completed a combined 47 sessions (Table 1). Participants were recruited from a past list of fMRI research participants to include individuals familiar with MRI scanning to reduce motion. Data from three sessions were not useable due to scan acquisition errors. Sessions were completed during evenings (18:00 - 21:00 start times) with a typical rate of 3-4 days between sessions (session information, acquisition errors, and timeline details provided in scan_sessions.csv on OSF).

**Table 1.** Sample Description.

| Participant | Sessions Completed | Age | Sex | Psychiatric Medications | Mean Framewise Displacement (mm) |
| --- | --- | --- | --- | --- | --- |
| sub-101 | 11 (9 useable) | 22 | Male | Sertraline | 0.067 |
| sub-103 | 12 (11 useable) | 23 | Female | Lisdexamfetamine | 0.233 |
| sub-104 | 12 | 24 | Male | None | 0.138 |
| sub-105 | 12 | 26 | Female | None | 0.153 |

The study design is summarized in Figure 1. In each session, participants completed the Positive and Negative Affect Schedule (PANAS) before scanning, Karolinska Sleepiness Scale (KSS) before and halfway through scanning, and two runs each of the MID and shared reward (SR) tasks. The order of the MID and SR tasks was counterbalanced across sessions. After the first runs of both tasks were completed, participants completed a positive mood induction task in the scanner. Mood induction involved reflecting on a positive memory that participants provided at the beginning of the session while listening to instrumental music intended to elicit positive mood (varied across sessions). New memories were provided by participants at each session and were presented as text in the scanner together with soothing music, with additional prompts encouraging reflection (e.g., “how did you feel in this moment”). T1w images were collected during the mood induction task. At the end of the session, participants completed a 10m resting-state scan or a neuromelanin scan, counterbalanced across sessions. Participants rated their momentary positive and negative mood before each acquisition (6 total observations).

**Figure 1.**
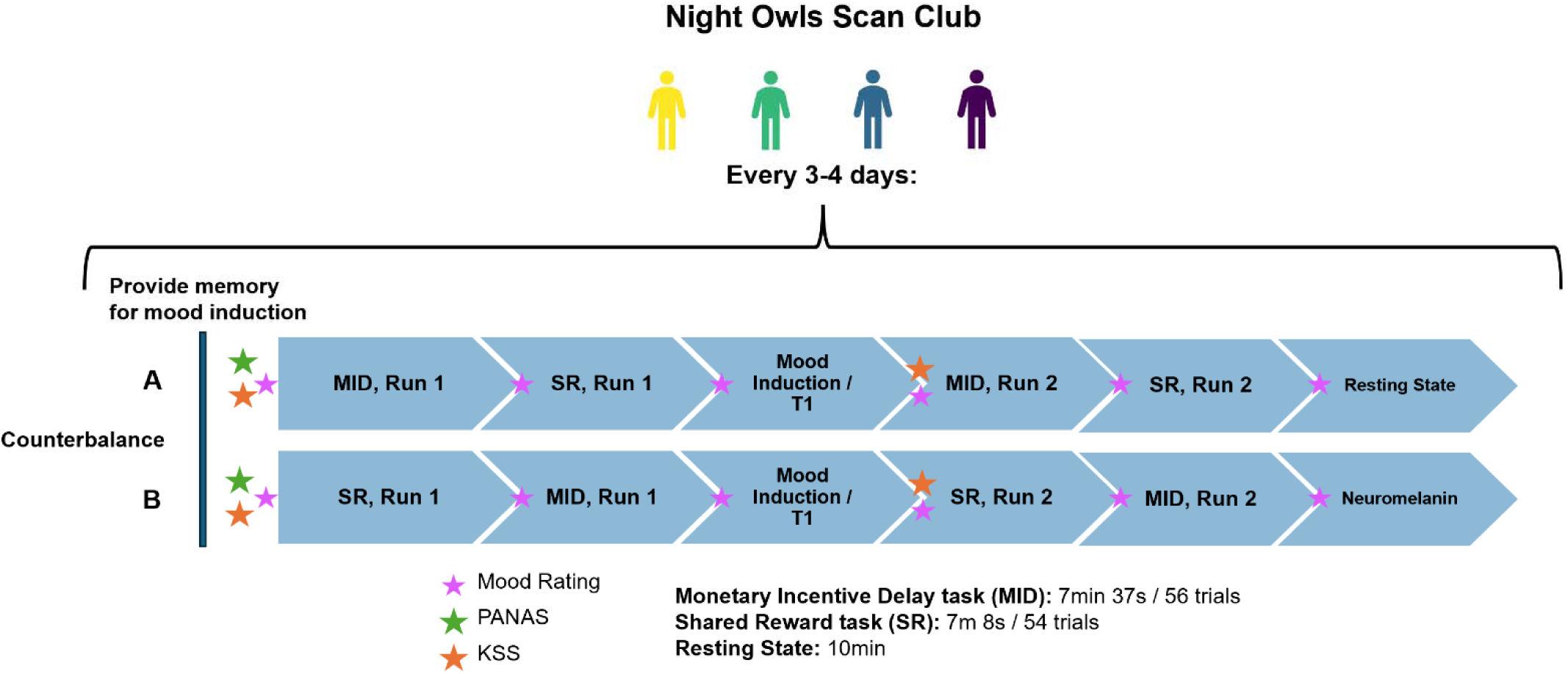
Study Design Summary. *Note*. MID = Monetary Incentive Delay task. SR = Shared Reward Task. PANAS = Positive and Negative Affect Schedule. KSS = Karolinska Sleepiness Scale.

### Measures

*PANAS*: The PANAS (Watson et al., 1988) is a 20-item self-report questionnaire that assesses positive and negative mood. Participants rated each item on a 1-5 point Likert scale reflecting the extent to which they experienced each item (e.g., “interested”, “distressed”) in the past four days or since their last scan, whichever was more recent. Positive and negative affect subscales were obtained from summing 10 respective items. We focused on positive affect in this study (PANAS-PA).

*KSS*: The KSS (Miley et al., 2016) is a self-report scale with a single item assessing sleepiness/alertness. Participants respond to a 1-9 Likert scale ranging from 1 “extremely alert” to 9 “extremely sleepy, can’t stay awake.” We reverse-coded scores such that higher scores indicate higher alertness.

*Caffeine Intake*: Participants indicated yes or no to if they had consumed caffeine in the four hours prior to the start of the session.

*Mood Ratings*: Participants rated their momentary positive and negative mood (“To what extent are you experiencing POSITIVE/NEGATIVE emotions RIGHT NOW?”) in the scanner on a 100-point continuous visual scale using a button box (10- and 1- point change intervals). As positive and negative mood ratings were highly anti-correlated, we only examined positive mood ratings here.

### Neuroimaging Acquisition

We collected anatomical and functional images using a 3.0 Tesla Siemens PRISMA MRI scanner and a 20-channel head coil at Temple University. T1w images were collected using a high-resolution magnetization-prepared rapid acquisition gradient echo (MPRAGE) sequence with in-plane acceleration (GRAPPA=2) with the following parameters: field of view (FOV) = 224 mm; matrix = 224×224; 1 mm^3^ isotropic resolution; 192 sagittal slices; repetition time (TR) = 2400ms; echo time (TE) = 2.17ms; flip angle = 8°. Multi-echo fMRI data were collected using a gradient echo-planar imaging (EPI) with four echoes (13.8, 31.54, 49.28, 67.02 ms) and simultaneous multislice (multiband factor = 3) with in-plane acceleration (GRAPPA = 2) and partial Fourier set to 7/8. This multi-echo sequence was collected with the following parameters: matrix = 80×80; 2.7 mm^3^ isotropic resolution; 10% gap between slices; 51 axial slices; TR = 1615ms; 50° flip angle. We collected both magnitude and phase components of the functional images, which facilitate denoising and undistortion approaches. Neuromelanin sensitive MRI images (not examined here) were also collected in counterbalance B (Figure 1) using a 2D gradient response echo sequence with magnetization transfer contrast.

### fMRI Tasks

Participants completed a version of the MID task (Knutson et al., 2000), an instrumental reward task where participants are rewarded based on performance to a timed button response (Supplementary Figure S1). To maximize the number of reward trials, we did not include a punishment condition. Participants were instructed to press their index finger button as soon as they see an image of a white square (“target”) in two different conditions. In neutral trials, the white square was preceded by a blue circle (“cue”) and participants were told they could not win or lose money but to still respond as quickly as possible. If participants responded in the allotted time, they were shown an outcome screen with a green check indicating successful response and black text stating “Trial earnings: $0.” In reward trials, the white square was preceded by a green circle and participants were told they could win $3. Successful responses were followed by an outcome screen of a green check and green text stating “Trial earnings: $3”. In both conditions, if participants failed to respond quickly enough, they were shown a black line and black text stating “Trial earnings: $0”. Cue screens were shown for 750ms, the interstimulus interval was jittered between 1500-3000ms, intertrial intervals (ITIs) were jittered between 2000-7000ms, target times were dynamically changed to maintain a 66% successful response rate, and outcome screens were shown for 1000ms. Each run contained 283 frames and 56 trials.

Participants also completed the Shared Reward task (Fareri et al., 2012), a card guessing task with reward and punishment in two different social contexts (Supplementary Figure S2). Participants were instructed that all cards for the task had an integer value ranging between 1 and 9, and their task on each trial was to indicate whether they thought the presented card would have a value above or below 5.

Participants were presented with feedback (1000ms) on if they had made the correct guess (reward) or an incorrect guess (punishment). Some trials contained neutral feedback (i.e., the value of the card equaled 5). Participants played each trial with one of two potential partners: a stranger (an image of an unknown peer) or a computer.

Participants were told strangers were previous participants in the study and that they would have an opportunity to be the “stranger” for subsequent participants following their participation. For every “reward” decision, both the participant and their “partner” won $5 each, whereas for every “punishment” decision, both the participant and their partner lost $2.50 each. Participants were told random trials would be selected for final reimbursement calculation. No money was won or lost on trials where neutral outcomes were experienced. The decision phase lasted until a decision was entered or a maximum 2500ms, the outcome phase lasted 1000ms, interstimulus intervals were jittered between 850-2550ms and trial-wise adjusted to account for decision phase timing, and ITIs were jittered between 1000-4000ms. Each run contained 265 frames and 54 trials. Outcomes were rigged such that each run summed to 22 win trials, 22 loss trials, and 10 neutral trials (order randomized across sessions).

An additional 10min (372 frames) of eye-open resting state scans (not examined here) were collected in counterbalance A (Figure 1). Participants were instructed to look at a crosshair and stay awake.

### Neuroimaging Data Preprocessing

Preprocessing was performed using fmriprep 24.1.1 (Esteban et al., 2019), which is based on Nipype 1.7.0 (Gorgolewski et al., 2011; RRID:SCR_002502). We used Warpkit (Van et al., 2023) to generate fieldmaps from multi-echo data for distortion correction and then used fmriprep in a two-stage approach. First, we passed all sessions of each subject through fmriprep with --anat-only and --longitudinal flags to generate a single T1w image per participant for consistency across sessions (Reuter et al., 2012). Second, the functional fmriprep pipeline was used for each session separately, with the subject-level T1w image as an existing derivative. BOLD runs were resampled onto MNI152NLin6Asym space. Full fmriprep preprocessing details are provided in the supplement. We used component-based and motion-based confounds for nuisance regression, as generated by fmriprep (see supplement), and high-pass filtering (128s cut-off) by using a set of discrete cosine basis functions. We also generated confound regressors identified by TEDANA as being artifactual and unrelated to true variation in BOLD signal (DuPre et al., 2021). To ensure identical smoothness across sessions and acquisitions, we used 3dBlurToFWHM with a smoothing kernel of 5mm. Finally, each run was grand-mean intensity normalized using a single multiplicative factor.

### fMRI Analysis and Moderating Factors

Neuroimaging analyses used FSL version 6.0.7 (Jenkinson et al., 2012; Smith et al., 2004). Each run of data was analyzed using a GLM with local autocorrelation (FILM). MID models included regressors for anticipation (cue onset through duration of due and interstimulus interval) and outcome (outcome onset through outcome duration) for reward and neutral conditions. SR models included regressors for the decision phase (decision onset through decision duration) and outcome phase (outcome onset through outcome duration) for each partner, with outcome regressors also split between gain, loss, and neutral.

For main analyses, we examined the Reward Anticipation > Neutral Anticipation contrast in the MID task and the Reward Outcome > Punishment Outcome (aggregated across partners) contrasts in the SR task. All models were estimated at an individual-level (L1) because the small sample does not permit population-level generalization.

We measured the reward response using anatomical univariate and canonical multivariate indices. As a univariate index, we extracted the mean Z score from a binary mask of the ventral striatum (VS) derived from the Oxford-GSK-Imanova striatal structural atlas (https://web.mit.edu/fsl_v5.0.10/fsl/doc/wiki/Atlases%282f%29striatumstruc.html;Tziortzi et al., 2011). As a multivariate index, we correlated unthresholded images with a previously validated multivariate brain signature of reward processing (https://neurovault.org/images/775976/; Speer et al., 2023).

### Trial-Level Models

To examine run-level reliability of reward responses, we estimated single-trial models using the Least Squares Single (LSS) approach, which helps alleviate collinearity across estimates for short inter-trial interval design (Mumford et al., 2012, 2014; Turner et al., 2012). LSS general linear models were estimated on each run with two regressors: one modelling event of interest in the specified trial (anticipation phase for MID, outcome phase for SR) and the other modelling all other trials’ events of interest in the run. We obtained one parameter estimate from the trial-specific regressor. This process was repeated for all trials. Consistent with other analyses, we also included the full set of the event regressors of non-interest in a run (MID outcome and SR decision) with their onsets and durations, as well as nuisance regressors from fmriprep and TEDANA.

### Moderating Factors

Finally, we examined how several methodological and analytical decisions influenced results. We specifically compared results using different combinations of the following decisions:

*Anatomical mask activation vs. similarity to canonical reward signature*: We compared results using the ventral striatal mask vs. a canonical brain reward signature, reflecting a univariate vs. multivariate operationalization of the reward response.

*Single-echo vs. multi-echo sequencing and denoising*: We compared results using optimally combined echoes vs. echo-2, following prior work (Giubergia et al., 2025; Heunis et al., 2021). For the optimally combined multi-echo data, we also examined the effect of including nuisance confounds estimated by TEDANA.

### Split-Half Reliability

Task responses are typically estimated across all trials in a run. However, variable responses across individual trials may limit validity of a run-averaged effect. We estimated the split-half reliability of trial-level BOLD responses, analogous to estimating internal consistency of items in a self-report scale. Due to the small sample size but larger number of functional runs, we estimated the split-half reliability across runs (i.e, across runs, correlation between halves of trial-level reward responses). We estimated the split-half reliability separately for pre- and post-mood induction runs using 1000 random equal trial splits for each, and then averaged reliability estimates together.

### Test-Retest Reliability

To examine the test-retest reliability of reward responses, we estimated multilevel models using the lme4 package (Bates et al., 2015) in R (R Core Team, 2022) and assessed ICC with the performance package (Lüdecke et al., 2021). For each contrast, we estimated the test-retest reliability across all pre-mood induction and post-mood induction runs (i.e., separate models for pre- and post-induction runs, ICCs then averaged together), and all session-level models (i.e., both runs included in same model). We focused on adjusted ICC estimates that account for fixed effects of session (e.g., detrending for habituation) and mean framewise displacement, as estimated by mriqc (Esteban et al., 2017).

### Intraindividual Associations with Mood and Alertness

Finally, we examined intraindividual associations between the reward response and mood and alertness. We used multilevel models with random effects of session (factorized) for all models, and random effects of subject for sample-pooled models. All variables were within-person standardized.

We first examined the effect of habituation and mood induction by estimating the effects of session and run, respectively, on VS activation and BRS correlation. We estimated a pooled model across the sample as well as separate idiographic (N = 1) models. We then examined intraindividual effects of run-level mood and alertness ratings and session-level PANAS-PA scores on reward responses in idiographic models. With limited generalizability of inferences due to the sample size, we focused on variance explained and correlation with mood ratings in idiographic models, rather than statistical significance.

## Results

### Mood and Alertness Ratings

Across participants, mood ratings immediately before and after the mood induction significantly increased by about 10 points (b = 10.4. SE = 1.9, p < .001). This effect was significant in idiographic models for participants 103, 104, and 105, but not 101 (Figure 2a). Consistent with general declines in mood observed in psychological research broadly (Jangraw et al., 2023), mood decreased in between other observations (i.e., outside of the mood induction; Figure 2b).

**Figure 2.**
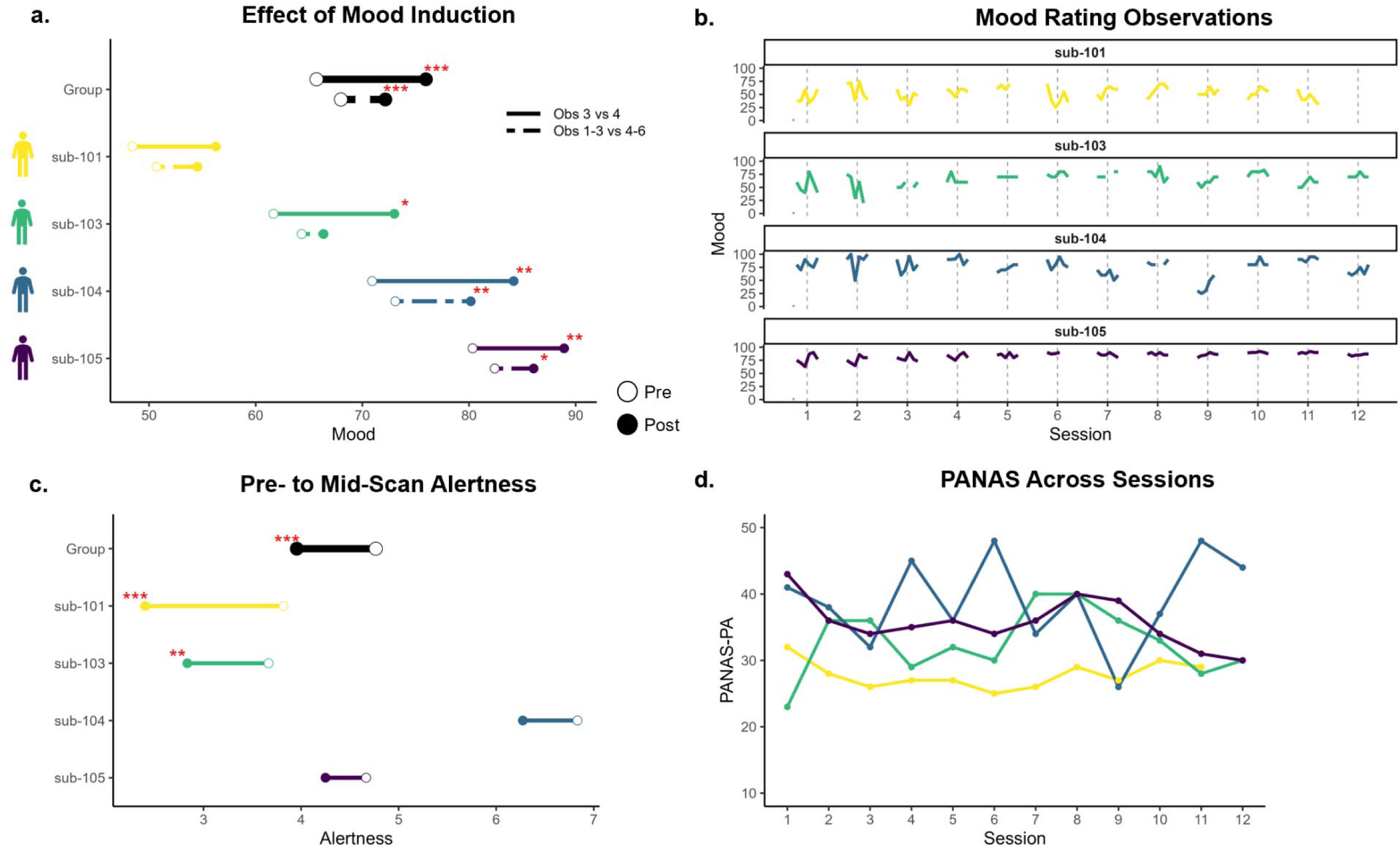
Mood and Alertness Ratings. *Note*. **a.** Effect of mood induction on self-reported mood ratings, comparing observations immediately before and after induction (solid line) and average of all ratings before and after induction (dashed line). **b.** Mood ratings across observations across sessions. **c.** Karolinska Sleepiness Scale (KSS) ratings before fMRI scan compared to halfway through fMRI scan (after mood induction). **d.** Positive and Negative Affect Schedule (PANAS) – Positive Affect (PA) subscale ratings across sessions.

Alertness significantly decreased across participants from the pre-scan rating to the mid-scan rating about halfway through the session (b = -0.80, SE = 0.14, p < .001). This effect was significant in idiographic models for participants 101 and 103, but not 104 or 105 (Figure 2c).

PANAS-PA ratings were relatively stable for participants 101 and 105, with more variability for participants 103 and 104 (Figure 2d). With a 10-50 score range, standard deviations were 2.0 for participant 101, 5.1 for participant 103, 6.6 for participant 104, and 3.7 for participant 105.

### Reward Response Split-Half Reliability

Single-trial models showed overall moderate internal consistency that was mostly consistent across participants (Figure 3ab). Split-half reliability tended to be slightly higher for VS activation relative to BRS correlation, but we did not find systematic differences between echo-2 vs. optimally combined multi-echo sequences, or base confounds vs. inclusion of TEDANA confounds. In the MID reward anticipation trials, the average split-half reliability was 0.61 for VS activation and 0.58 for BRS correlation. In SR reward outcome trials, the average split-half reliability was 0.48 for VS activation and 0.45 for BRS correlation.

**Figure 3.**
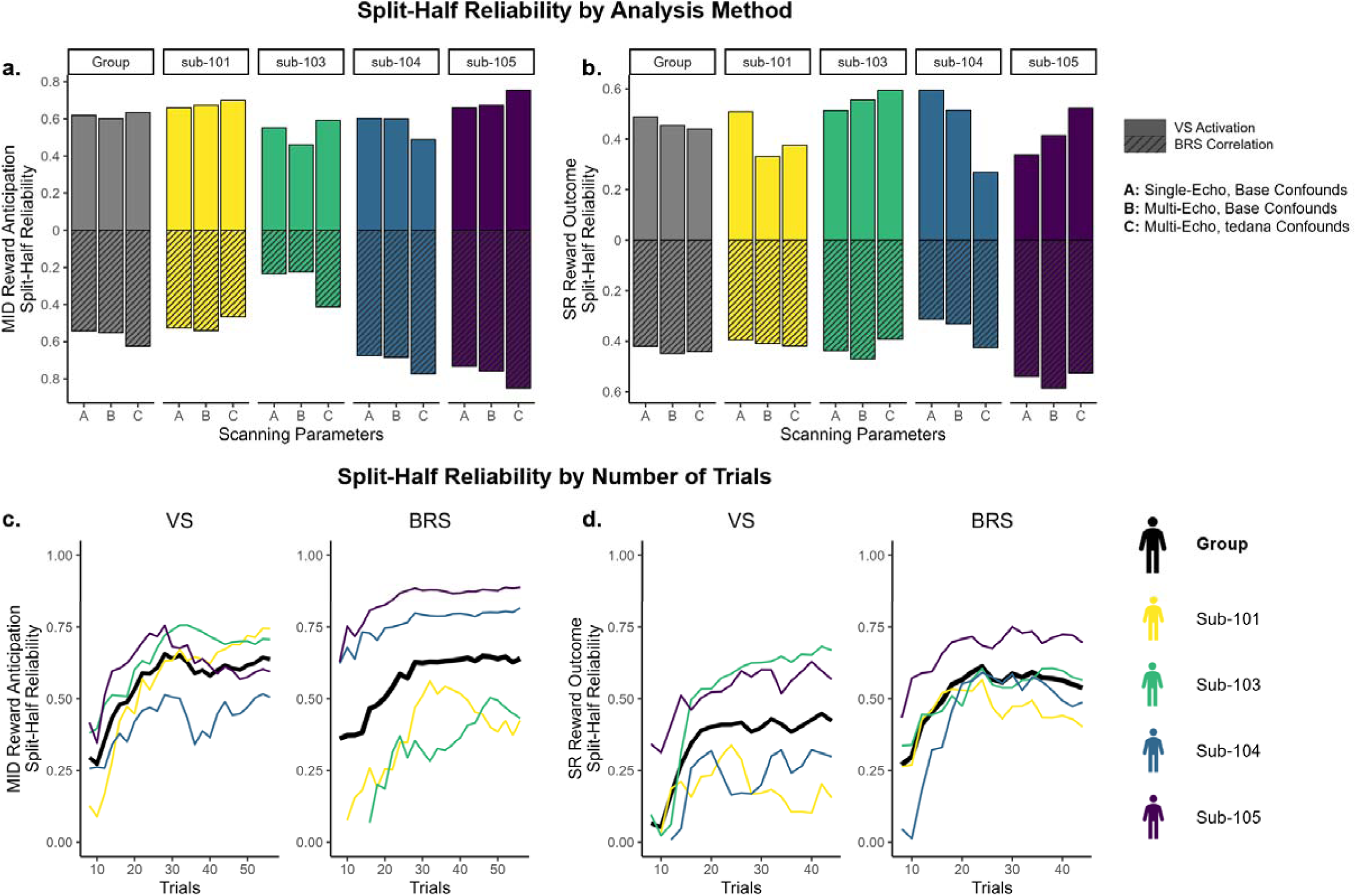
Split-Half Reliability of Single-Trial Models. *Note.* MID = Monetary Incentive Delay task. SR = Shared Reward task. **a,b.** Split-half reliability in models estimated with single-echo vs. multi-echo scanning sequences and confounds, and anatomical ventral striatum (VS) mask vs. multivariate brain reward signature (BRS) mask. **c,d.** Split-half reliability in session-wise models (averaged across runs) estimated with different number of trials, ranging from 8 to maximum.

We also examined how split-half reliability was moderated by the number of trials (Figure 3cd). Using the optimally combined multi-echo sequences and TEDANA confounds, we concatenated trials within sessions and regressed out the effect of run. We then selected increasing numbers of trials from 8 to the maximum number of trials (56 for MID reward anticipation, 44 for SR reward outcome) and repeated split-half reliability analyses. Split-half reliability increased with the number of trials but plateaued around 25 reward-condition trials for VS activation and 20 reward-condition trials for BRS correlation.

### Reward Response Test-retest Reliability

Test-retest reliability was low, ranging from .07 to .15 for MID and near 0 to 0.36 for SR (Figure 4). ICC estimates were generally higher for BRS correlation than VS activation, but there were no systematic differences in single-echo vs. optimally combined multi-echo sequences or confound regression. Test-retest reliability was higher in BRS correlation relative to VS activation, especially with SR reward outcome (ICC estimates around 0.35). However, due to the small sample size, multilevel models did not converge for VS activation estimates in the SR task for (1) run-level models using multi-echo sequences and confounds, (2) run-level models using single-echo sequences and base confounds, or (3) session-level models with multi-echo sequences and confounds (Figure 4). Consistent with low test-retest reliability, reward responses were highly variable across sessions (Supplementary Figure S3).

**Figure 4.**
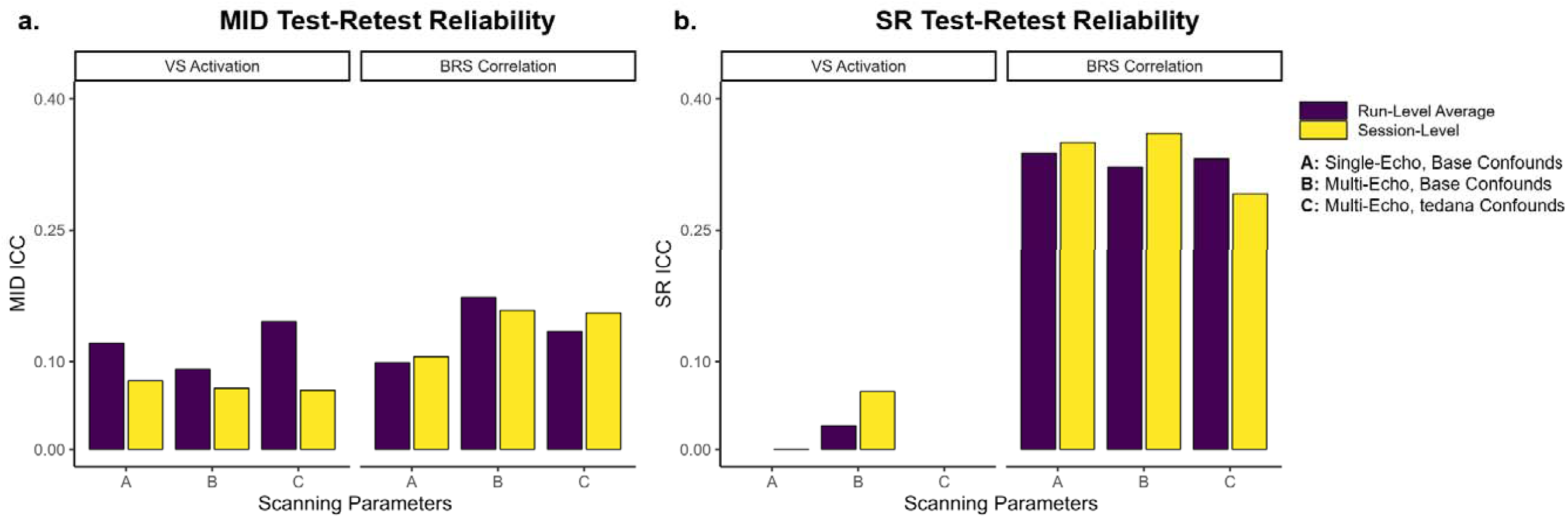
Test-Retest Reliability. *Note.* MID = Monetary Incentive Delay task. SR = Shared Reward task. VS = Ventral Striatum. BRS = Brain Reward Signature. Run-level average reflects the average ICC between separate multilevel models for pre-mood induction runs and post-mood induction runs. Session-level reflects ICC from multilevel models that included effects of session and each run. Models did not converge for S VS activation run-level models for scanning sequences A or C, or session-level models for scanning sequence C.

### Intraindividual Changes and Behavioral Associations

We first examined intraindividual effects of session (i.e., habituation) and run (i.e., mood induction) on the reward response. Across the sample, there were no significant effects of session or run on VS activation or BRS correlation in MID anticipation or SR outcome contrasts (Supplementary Table S1). Idiographic responses before and after the mood induction are shown in Figure 5. For the MID task, participants had small but consistent increases in VS activation and BRS correlation after the mood induction. In the SR task, results were more variable across participants and masks.

**Figure 5.**
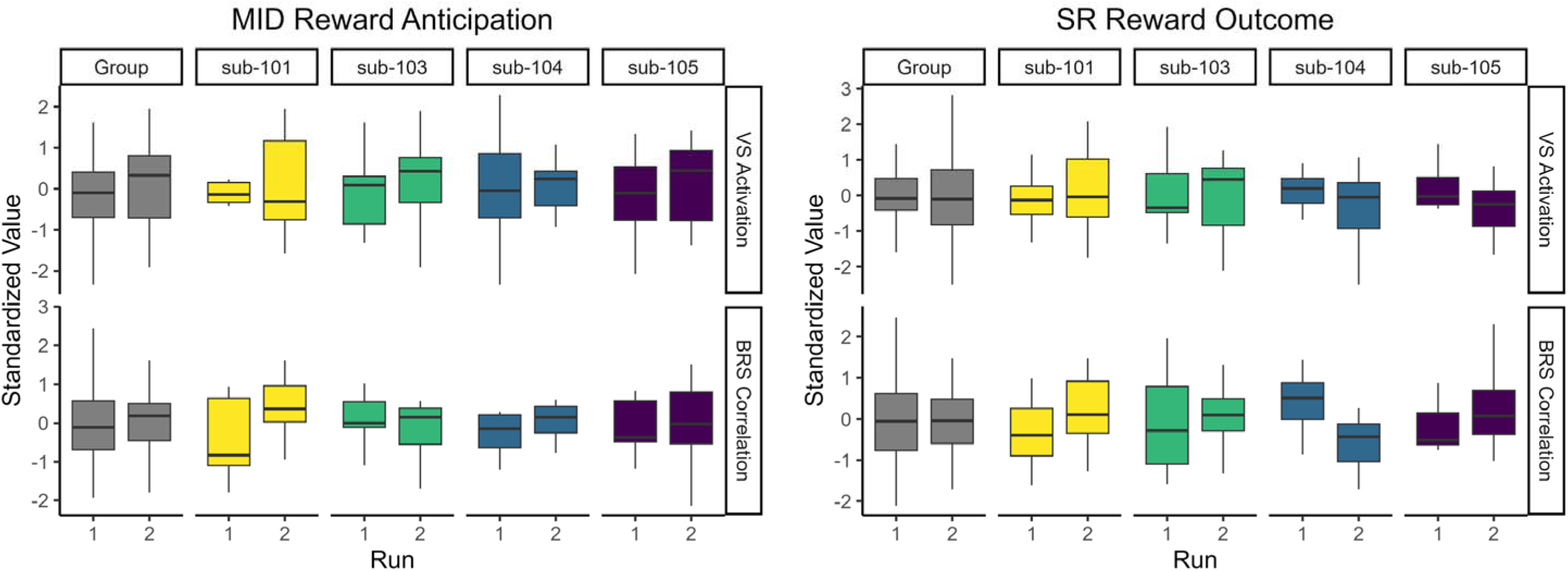
Reward Responses Pre- and Post-Mood Induction. *Note*. VS = Ventral Striatum. BRS = Brain Reward Signature. Run 1 and run 2 indicate reward response pre- and post-mood induction, respectively.

We then examined intraindividual associations between reward responses and mood and alertness ratings in idiographic models. On average, mood and alertness explained 10% variance of MID VS activation, 20% variance of MID BRS correlation, 9% variance of SR VS activation, and 8% variance of SR BRS correlation. BRS correlation in MID anticipation had the most variance explained by intraindividual mood and alertness, including 31% for sub-103. Table 2 provides marginal and semi-partial effects of mood ratings, PANAS-PA, and alertness (displayed in Supplementary Figure S4).

**Table 2.** Intraindividual Variance Explained by Mood and Alertness.

|  | **Sub-101** |  | **Sub-103** |  | **Sub-104** |  | **Sub-105** |  |
| --- | --- | --- | --- | --- | --- | --- | --- | --- |
| **MID Anticipation** | VS | BRS | VS | BRS | VS | BRS | VS | BRS |
| **Marginal R<sup>2</sup>** | .225 | .150 | .087 | .309 | .067 | .153 | .016 | .186 |
| **Semi-Partial R<sup>2</sup>** |  |  |  |  |  |  |  |  |
| Mood Ratings | .138 | .015 | .036 | .166 | .001 | .079 | .012 | 0 |
| PANAS-PA | .093 | .139 | .022 | .189 | .030 | .038 | 0 | .035 |
| KSS Alertness | .001 | .036 | .002 | .108 | .005 | .083 | .002 | .146 |
| **SR Outcome** |  |  |  |  |  |  |  |  |
| **Marginal R<sup>2</sup>** | .136 | .114 | .064 | .020 | .085 | .037 | .068 | .154 |
| **Semi-Partial R<sup>2</sup>** |  |  |  |  |  |  |  |  |
| Mood Ratings | .054 | 0 | .010 | 0 | 0 | 0 | .065 | 0 |
| PANAS-PA | .012 | .103 | 0 | 0 | .026 | 0 | .052 | .022 |
| KSS Alertness | 0 | .002 | 0 | 0 | 0 | 0 | .054 | .120 |

*Note.* Marginal R<sup>2</sup> reflects variance explained by fixed effects. Semi-partial R<sup>2</sup> reflects unique contribution by each predictor, beyond variance explained by other predictors.

## Discussion

Neuroimaging research mostly relies on between-person comparisons, but tests of individual differences are limited by low test-retest reliability and cannot inform within-person processes. This study examined if low test-retest reliability of the BOLD reward response is explained by measurement error and long scanning intervals or reflects true within-person change with behavioral associations in mood and alertness. We collected and openly released a novel intensively sampled fMRI study featuring within-session mood manipulations - the Night Owls Scan Club. We also examined how precision imaging methodologies (e.g., multi-echo sequencing and confound regression, multivariate reward signatures) affected measurement properties.

Despite the short retest intervals, test-retest reliability of the reward response was very low, ranging from near 0 to 0.15 for an anatomical ventral striatal mask and 0.10 to 0.35 for a multivariate reward signature. Furthermore, low test-retest reliability was not explained by measurement error, as individual runs showed acceptable split-half reliability and participants demonstrated relatively low motion. Instead, results were more indicative of within-person change. While there was not a significant sample-wide effect of the mood induction, reward anticipation responses showed small but consistent increases after the mood induction in idiographic models. Moreover, intraindividual mood and alertness ratings generally explained 10-20%, and as much as 31% of the variance in within-person fluctuations in the reward response, with mood explaining relatively more variance. Together, results suggest that the BOLD task responses are better explained by within-person variance, not between-person variance. Considering similar findings in a different emotion processing task (Flournoy et al., 2024) and generally low test-retest reliability (Elliott et al., 2020) and stability (Gratton et al., 2018) of task-based activations, we suggest that task-based fMRI research should shift its traditional focus on between-person comparisons to within-person study. With the scarcity of within-person study designs, there is substantial opportunity for novel insights into intraindividual functioning that can inform more mechanistic hypotheses. To help reach this potential, we provide open data from this novel precision fMRI study with intensive sampling and experimental manipulations.

### Low Test-Retest Reliability is Not Explained by Measurement Error

Our findings extend prior research showing that the BOLD reward response is not a stable, trait-like marker within individuals (Demidenko et al., 2024; Elliott et al., 2020). Data here were acquired days apart with multiband, multi-echo scanning sequences, have relatively low motion, and showed acceptable split-half reliability.

Despite the properties, most notably a retest interval that is magnitudes shorter than most fMRI studies, test-retest reliability was very low. Test-retest reliability was also not improved by multi-echo sequencing or denoising. Moreover, considering the noise present within individual task trials, we believe that split-half reliability results of moderate-to-strong trial-level correlations suggest that individual runs are reliable measurements of the reward response, further dissuading concerns that low test-retest reliability is due to measurement noise. Together, converging findings from this intensive sampling study, a meta-analysis (Elliott et al., 2020), and a multiverse analysis of different analytical approaches (Demidenko et al., 2024) strongly suggest that the BOLD reward response is not a stable trait-like marker. We thus echo Elliott et al. (2020) that the reward response, like most task-based BOLD responses, is of little utility for tests of individual differences, especially without very large samples.

### Reward Responses Reflect Intraindividual Fluctuations in Mood and Alertness

In contrast to limited utility for between-person comparisons, results suggest high potential for within-person research. To enable intraindividual investigation, we used a novel intensive fMRI sampling design and behavioral manipulation. The mood induction successfully increased positive mood by about 10 points on a 100-point scale. This increase contrasts observed decreases in mood between other observations without the mood induction, robust evidence that mood generally declines across a research study (Jangraw et al., 2023), and general expectations that positive mood would not increase after being in an fMRI scanner for more than 30 minutes. The increase in mood is also opposite to decreases in alertness within sessions. As alertness is associated with functional connectivity (Laumann et al., 2017; Tagliazucchi & Laufs, 2014), future precision imaging research including long acquisitions should measure and account for changes in alertness across acquisitions.

We were specifically interested in how the reward response changed pre-to-post mood induction and was associated with intraindividual fluctuations in mood and alertness. There were no sample-wide significant changes in the reward response after the mood induction. However, in idiographic models, reward anticipation contrasts in the MID task showed systematic increases for all participants in both univariate and multivariate indices, except the univariate response for sub-101. In contrast, mood induction-related changes in reward outcome contrasts in the SR task were variable across the sample, with the aggregated effect reflecting essentially no change. While caution is warranted based on the sample size and relatively high standard errors, increased response to reward anticipation specifically (not outcome) following a mood induction is consistent with prior work (Young & Nusslock, 2016). We extended this finding to show the effect may be consistent across numerous sessions. These findings suggest that increasing an individual’s mood increases their neural response to reward anticipation. We also found that mood ratings, PANAS-PA scores, and alertness scores explained about 10-20% of idiographic variance in reward responses across runs and sessions, including as much as 31% of the variance for one participant. While this is an appreciable effect for fMRI, we note one potential limiting factor here in that mood ratings and PANAS-PA scores tended to have little within-person variation across sessions. Overall, results here provide novel evidence of relationships between neural reward responses and mood and alertness at a within-person, rather than between-person level. Further intraindividual study of mood and reward functioning may inform how brain functioning changes with the onset or treatment of low mood in clinical disorders (e.g., anhedonia).

### Optimizing Task-Based Split-Half Reliability

Relative to rest-retest reliability, split-half reliability of trials across runs was notably higher. While split-half reliability near 0.60 would be low for a self-report scale, we believe that considering the noise present within individual trials, these results provide confidence that contrast estimates from a single run reliably reflect responses across that run. Moreover, split-half reliability increased with the number of trials, analogous to findings that reliability of resting state connectivity networks increases with more data (Gordon et al., 2017; Ooi et al., 2025). Increases in split-half reliability plateaued around 20-25 *reward* trials. Since control condition trials (e.g., neutral or loss) are also needed for contrasts, nearly 50 trials may be recommended for more reliable task response measurement. Additionally, using jittered ITIs further improves BOLD measurement (Miezin et al., 2000; Ollinger et al., 2001). However, many task designs include 10-20 trials per run and no ITI, such as the frequently examined Adolescent Brain Cognitive Development study (Casey et al., 2018), potentially limiting reliability.

Following recent advances in resting state connectivity research (Ooi et al., 2025), task-based fMRI research should empirically examine how to optimally allocate resources between task length (number of trials and ITI) and sample size. Noting that split-half reliability plateaued around 25 *reward* trials in this study and previous findings that nearly 60 trials were needed to maximize test-retest reliability in a pain processing task (Han et al., 2022), it is likely that substantially more resources should be allocated to individual task runs than what is typically provided.

### Precision Imaging Methodological Factors

We were interested in how several methodological decisions relevant to precision imaging altered results. First, we compared optimally combined multi-echo sequences and denoising based on TEDANA with single-echo (echo-2) sequences and denoising solely based on fmriprep confounds. In contrast to expectations, we observed no benefit of multi-echo sequences or denoising. Previous work has found that multi-echo sequences generally show improved signal-to-noise and reliability (Giubergia et al., 2025; Gonzalez-Castillo et al., 2016; Heunis et al., 2021). One potential explanation here could be that multi-echo sequences have been shown to be less sensitive to task events (Giubergia et al., 2025; Gonzalez-Castillo et al., 2016). More research is needed to understand the costs and benefits of multi-echo sequencing in task-based designs, especially for activation in subcortical regions such as the VS.

Second, we compared results using a univariate mask of VS activation to a multivariate canonical reward signature (BRS). Relative to VS activation, contrast map correlations with the BRS showed higher test-retest reliability, reached their peak split-half reliability in fewer trials, and were more strongly associated with mood and alertness in the MID task. These findings extend prior evidence of higher test-retest reliability in multivariate signatures (e.g., Kragel et al., 2021) to suggest that multivariate signatures are also more internally consistent and sensitive to intraindividual change.

Thus, multivariate signatures appear to be a promising improvement to univariate study and present an exciting direction for task-based fMRI research.

### Limitations

This study has several limitations that contextualize results and that future research should address. Similar to other precision imaging studies, the main limitation is the very small sample size (N=4). While the small sample size enabled us to focus resources on maximizing reliability of within-person associations for each individual, the generalizability of those within-person relationships is low – a common criticism of idiographic research. The small sample size also prevents between-person comparison of mood and reward responses. Further limiting generalizability, our sample contained all White, educated, young adults. Due to time and resource requirements, precision imaging research has so far largely failed to recruit diverse samples, with other studies also relying on researchers serving dual roles as participants. To avoid intensively sampled neuroimaging being mostly based on a single minority demographic, future research must explicitly focus on recruiting more diverse samples. Another limitation is the risk that participants habituated to reward stimuli with repeated exposure, a challenge for intensive task-based designs. While we did not observe systematic decreases in the reward response across sessions here, future work should consider developing task variations that alter stimuli to reduce habituation but maintain task fidelity. Finally, while mood significantly increased following the mood induction, changes were relatively small, potentially limiting effects on fMRI associations. PANAS- PA and mean momentary mood ratings also had little variation across sessions, limiting power for within-person associations. Most self-report scales are designed and validated to maximize cross-sectional reliability and may thus be less appropriate for within-person study. Future research may consider combining intensive neuroimaging with ecological momentary assessment and passive monitoring methods to increase within-person variance.

### Conclusions and Future Directions: Intraindividual Neuroimaging

We collected and publicly released NOSC - a novel intensively sampled fMRI study focused on reward functioning, mood, and precision imaging methodology.

Findings of low test-retest reliability suggest that the reward-related task response is of limited utility for between-person comparisons. In contrast, we found evidence that positive mood manipulations may increase reward anticipation responses and that mood and alertness explain about 20% of intraindividual fluctuations in the reward response. Finally, we found that a multivariate reward signature was more reliable and sensitive to change than activation in a univariate striatal mask, but multi-echo sequencing or denoising had little effects. Overall, we present novel data and findings that underscore the need to transition from a reliance on between-person study to also examining intraindividual brain-behavior associations, especially for task-based fMRI. As many research questions with important implications are based on the causes and consequences of changes in brain functioning, there is substantial opportunity for novel insights through intensively sampled designs.

Moreover, this study highlights several key points for the future of intraindividual fMRI research. First and most importantly, within-person inferences cannot be made from between-person research (Mattoni, Fisher, et al., 2025). There is a large mismatch between the importance of within-person hypotheses (e.g., how changes in brain functioning lead to depression onset) and the scarcity of within-person study designs.

While between-person research can inform many important questions related to individual differences, there is substantial need for increased attention to intraindividual research. This study provides one example and a resource for future work to build on.

Second, task-based activations are particularly promising for intraindividual research. Precision imaging research has demonstrated that large-scale functional connectivity networks are largely stable given sufficient data (Gordon et al., 2017; Gratton et al., 2018). In contrast, task-based activations are less stable, more associated with state factors, and have low test-retest reliability (Elliott et al., 2020;

Flournoy et al., 2024; Gratton et al., 2018). Although they are related (Laumann et al., 2015), it is possible that functional connectivity approaches may be more appropriate for between-person study, while task-based activation is more appropriate for within-person study.

Third, for intraindividual research, the number of observations (T) is as important, if not more so, than the sample size (N). Investigators must carefully balance N and T, depending on their goals (Gell et al., 2024). As we have very little current knowledge of intraindividual brain-behavior relationships, idiographic study is an apt starting point before gathering larger samples to assess generalizability. In this study, collecting 12 sessions of data, with each having an experimental manipulation, provided up to 24 observations per task per person. This framework highlights the feasibility of intensive intraindividual assessment using fMRI that can easily be extended into other areas of interest.

Fourth, time is a critical consideration in within-person research and the appropriate interval length between assessments is an unresolved question.

Longitudinal fMRI research mostly examines scans that are months or years apart, limiting ability to assess within-person brain-behavior relationships. While more intensive intervals are needed, the most appropriate temporal scale is unclear. We measured mood before entering the scanner (PANAS) as well as seconds to minutes before starting scans (mood ratings). However, there may be distinct effects when examined contemporaneously or with time lags ranging from milliseconds to days. This is an important empirical question to address as intensive longitudinal neuroimaging grows.

Finally, continuing precision imaging trends of open data (e.g., Gordon et al., 2017) will be fundamental to increasing accessibility and reproducibility of intraindividual research. As precision imaging research has so far largely been used for data aggregation to maximize reliability of between-person comparisons, there is current opportunity for within-person study using existing datasets (see Table 1 in Michon et al., 2022 for one list of prior works).

## CRediT Authorship Statement

**MM**: Conceptualization, Project Administration, Formal Analysis, Investigation, Writing – Original Draft, Visualization. **SW:** Formal Analysis, Project Administration, Writing – Review and Editing. **CJS:** Project Administration, Writing – Review and Editing**. TMO:** Conceptualization, Supervision, Writing – Review and Editing. **DVS:** Conceptualization, Formal Analysis, Writing – Review and Editing, Supervision, Project Administration, Funding Acquisition.

## Acknowledgements

We would like to thank the generous contributions of the participants, as well as research assistants Ashley Hawk, Jenelle Scholl, Jamie-Nicole Luistro, Ryan Gephart, and Avi Dachs, who made this study possible.

