## Supplementary Materials for "Precision Imaging for Intraindividual Investigation of the Reward Response"

**Supplementary Figure S1. Monetary Incentive Delay Task Schematic**

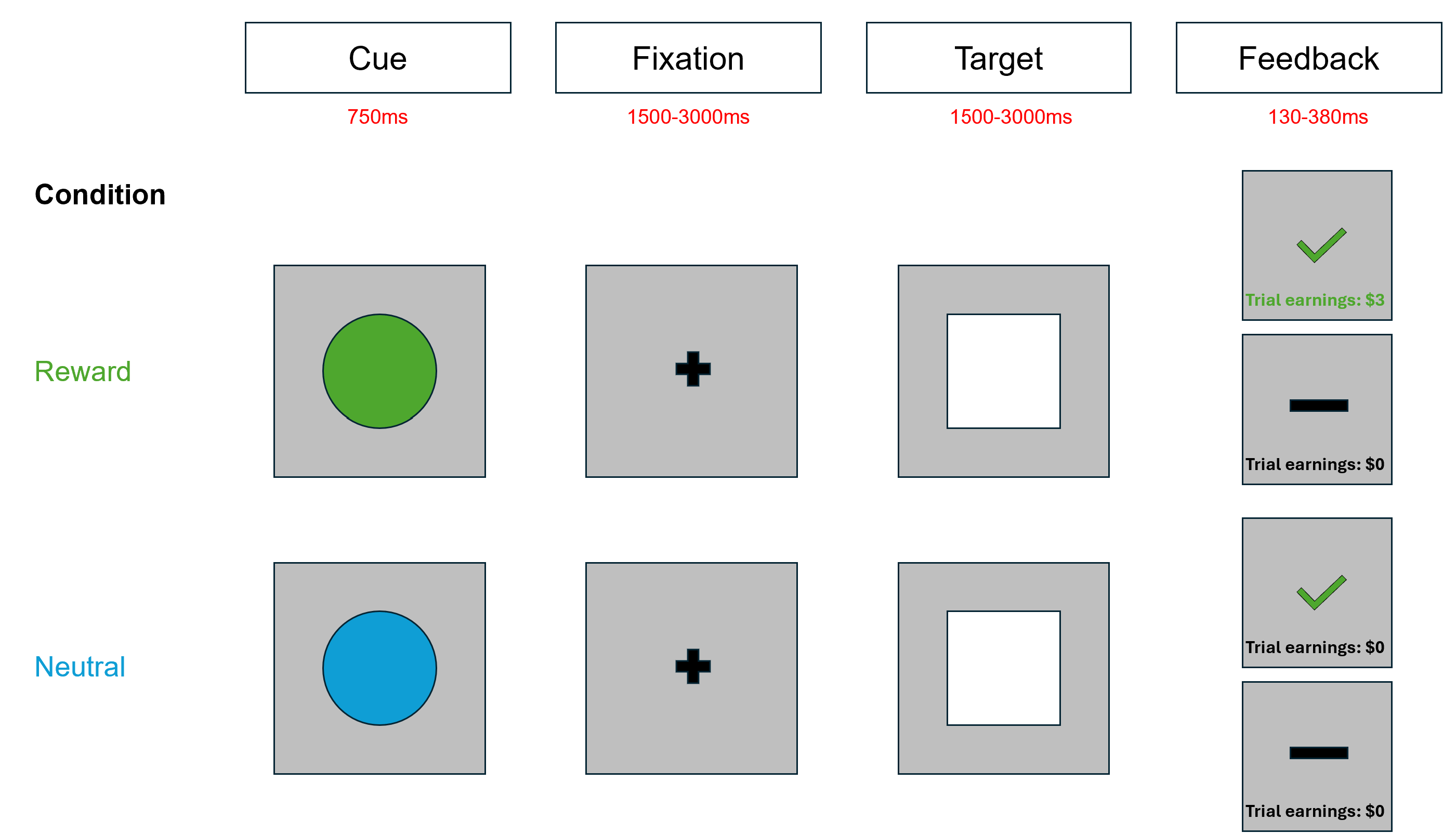

*Note*. ITIs jittered between 2000-7000ms before repeating trials.

**Supplementary Figure S2. Shared Reward Task Schematic**

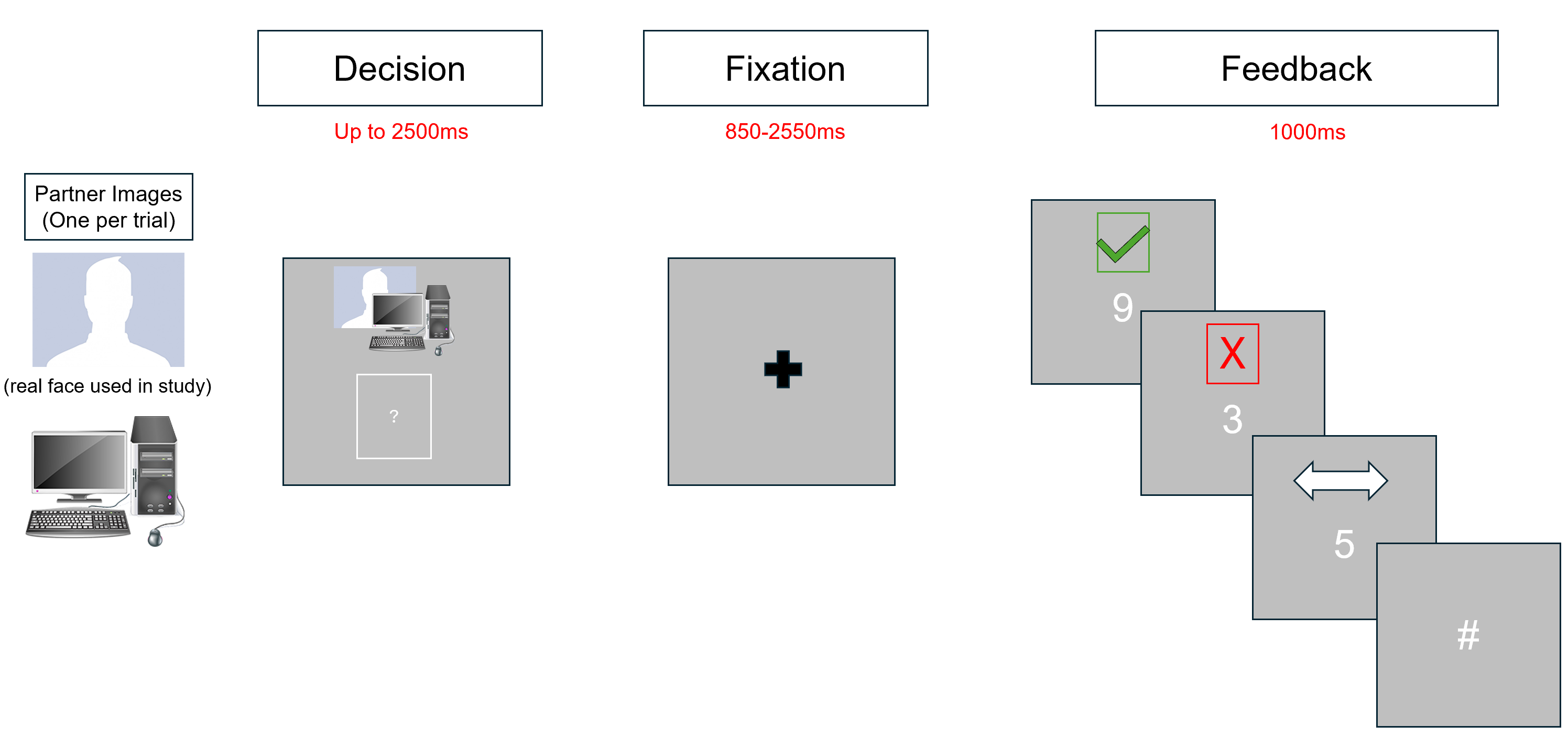

*Note.* ITIs jittered between 1000-4000ms before repeating trials. Pound sign appeared when participants failed to respond within 2500ms.

**Supplementary Methods: Neuroimaging Data Preprocessing**

Results included in this manuscript come from preprocessing performed using *fMRIPrep* 24.1.1 (Esteban et al. (2019); Esteban et al. (2018); RRID:SCR_016216), which is based on *Nipype* 1.8.6 (K. Gorgolewski et al. (2011); K. J. Gorgolewski et al. (2018); RRID:SCR_002502).

**Stage 1 Anatomical data preprocessing**

Each T1w image was corrected for intensity non-uniformity (INU) with N4BiasFieldCorrection (Tustison et al. 2010), distributed with ANTs 2.5.3 (Avants et al. 2008, RRID:SCR_004757). The T1w-reference was then skull-stripped with a *Nipype* implementation of the antsBrainExtraction.sh workflow (from ANTs), using OASIS30ANTs as target template. Brain tissue segmentation of cerebrospinal fluid (CSF), white-matter (WM) and gray-matter (GM) was performed on the brain-extracted T1w using fast (FSL (version unknown), RRID:SCR_002823, Zhang, Brady, and Smith 2001). An anatomical T1w-reference map was computed using mri_robust_template (FreeSurfer 7.3.2, Reuter, Rosas, and Fischl 2010). Volume-based spatial normalization to two standard spaces (MNI152NLin6Asym, MNI152NLin2009cAsym) was performed through nonlinear registration with antsRegistration (ANTs 2.5.3), using brain-extracted versions of both T1w reference and the T1w template. The following templates were selected for spatial normalization and accessed with *TemplateFlow* (24.2.0, Ciric et al. 2022): *FSL’s MNI ICBM 152 non-linear 6th Generation Asymmetric Average Brain Stereotaxic Registration Model* [Evans et al. (2012), RRID:SCR_002823; TemplateFlow ID: MNI152NLin6Asym], *ICBM 152 Nonlinear Asymmetrical template version 2009c* [Fonov et al. (2009), RRID:SCR_008796; TemplateFlow ID: MNI152NLin2009cAsym].

Many internal operations of *fMRIPrep* use *Nilearn* 0.10.4 (Abraham et al. 2014, RRID:SCR_001362), mostly within the functional processing workflow. For more details of the pipeline, see [the section corresponding to workflows in *fMRIPrep*’s documentation](https://fmriprep.readthedocs.io/en/latest/workflows.html).

**Preprocessing of B_0_ inhomogeneity mappings**

A total of 5 fieldmaps were found available within the input BIDS structure for each subject. A *B_0_* nonuniformity map (or *fieldmap*) was directly measured with an MRI scheme designed with that purpose such as SEI (Spiral-Echo Imaging).

**Stage 2 Anatomical data preprocessing**

T1-weighted (T1w) images were found within the input BIDS dataset. A preprocessed T1w image was provided as a precomputed input and used as T1w-reference throughout the workflow. A pre-computed brain mask was provided as input and used throughout the workflow. Precomputed discrete tissue segmentations were provided as inputs. Precomputed tissue probabiilty maps were provided as inputs.

**Functional data preprocessing**

For each of the 5 BOLD runs found per subject (across all tasks and sessions), the following preprocessing was performed. First, a reference volume was generated from the shortest echo of the BOLD run, using a custom methodology of *fMRIPrep*, for use in head motion correction. Head-motion parameters with respect to the BOLD reference (transformation matrices, and six corresponding rotation and translation parameters) are estimated before any spatiotemporal filtering using mcflirt (FSL , Jenkinson et al. 2002). The estimated *fieldmap* was then aligned with rigid-registration to the target EPI (echo-planar imaging) reference run. The field coefficients were mapped on to the reference EPI using the transform. The BOLD reference was then co-registered to the T1w reference using mri_coreg (FreeSurfer) followed by flirt (FSL , Jenkinson and Smith 2001) with the boundary-based registration (Greve and Fischl 2009) cost-function. Co-registration was configured with six degrees of freedom. Several confounding time-series were calculated based on the *preprocessed BOLD*: framewise displacement (FD), DVARS and three region-wise global signals. FD was computed using two formulations following Power (absolute sum of relative motions, Power et al. (2014)) and Jenkinson (relative root mean square displacement between affines, Jenkinson et al. (2002)). FD and DVARS are calculated for each functional run, both using their implementations in *Nipype* (following the definitions by Power et al. 2014). The three global signals are extracted within the CSF, the WM, and the whole-brain masks. Additionally, a set of physiological regressors were extracted to allow for component-based noise correction (*CompCor*, Behzadi et al. 2007). Principal components are estimated after high-pass filtering the *preprocessed BOLD* time-series (using a discrete cosine filter with 128s cut-off) for the two *CompCor* variants: temporal (tCompCor) and anatomical (aCompCor). tCompCor components are then calculated from the top 2% variable voxels within the brain mask. For aCompCor, three probabilistic masks (CSF, WM and combined CSF+WM) are generated in anatomical space. The implementation differs from that of Behzadi et al. in that instead of eroding the masks by 2 pixels on BOLD space, a mask of pixels that likely contain a volume fraction of GM is subtracted from the aCompCor masks. This mask is obtained by thresholding the corresponding partial volume map at 0.05, and it ensures components are not extracted from voxels containing a minimal fraction of GM. Finally, these masks are resampled into BOLD space and binarized by thresholding at 0.99 (as in the original implementation). Components are also calculated separately within the WM and CSF masks. For each CompCor decomposition, the *k* components with the largest singular values are retained, such that the retained components’ time series are sufficient to explain 50 percent of variance across the nuisance mask (CSF, WM, combined, or temporal). The remaining components are dropped from consideration. The head-motion estimates calculated in the correction step were also placed within the corresponding confounds file. The confound time series derived from head motion estimates and global signals were expanded with the inclusion of temporal derivatives and quadratic terms for each (Satterthwaite et al. 2013). Frames that exceeded a threshold of 0.5 mm FD or 1.5 standardized DVARS were annotated as motion outliers. Additional nuisance timeseries are calculated by means of principal components analysis of the signal found within a thin band (*crown*) of voxels around the edge of the brain, as proposed by (Patriat, Reynolds, and Birn 2017). All resamplings can be performed with *a single interpolation step* by composing all the pertinent transformations (i.e. head-motion transform matrices, susceptibility distortion correction when available, and co-registrations to anatomical and output spaces). Gridded (volumetric) resamplings were performed using nitransforms, configured with cubic B-spline interpolation.

Many internal operations of *fMRIPrep* use *Nilearn* 0.10.4 (Abraham et al. 2014, RRID:SCR_001362), mostly within the functional processing workflow. For more details of the pipeline, see [the section corresponding to workflows in *fMRIPrep*’s documentation](https://fmriprep.readthedocs.io/en/latest/workflows.html).

**Copyright Waiver**

The above boilerplate text was automatically generated by fMRIPrep with the express intention that users should copy and paste this text into their manuscripts *unchanged*. It is released under the [CC0](https://creativecommons.org/publicdomain/zero/1.0/) license.

**Confound Regressors**

A set of physiological regressors were extracted to allow for component-based noise correction (CompCor). These components are estimated after high-pass filtering; the preprocessed BOLD time-series (using a discrete cosine filter with 128s cut-off) for anatomical component correction (aCompCor). For aCompCor, three probabilistic masks (CSF, WM and combined CSF+WM) are generated in anatomical space. The implementation differs from that of Behzadi et al. in that instead of eroding the masks by 2 pixels on BOLD space, the aCompCor masks are subtracted from a mask of pixels that likely contain a volume fraction of GM. This mask is obtained by thresholding the corresponding partial volume map at 0.05, and it ensures components are not extracted from voxels containing a minimal fraction of GM. Finally, these masks are resampled into BOLD space and binarized by thresholding at 0.99 (as in the original implementation). Components are also calculated separately within the WM and CSF masks. For each CompCor decomposition, the k components with the largest singular values are retained, such that the retained components’ time series are sufficient to explain 50 percent of variance across the nuisance mask (CSF, WM, combined, or temporal). The remaining components are dropped from consideration. The head-motion estimates calculated in the correction step also were placed within the corresponding confounds file. All resamplings can be performed with a single interpolation step by composing all the pertinent transformations (i.e., head-motion transform matrices, susceptibility distortion correction when available, and co-registrations to anatomical and output spaces). Gridded (volumetric) resamplings were performed using ‘antsApplyTransforms’ (ANTs), configured with Lanczos interpolation to minimize the smoothing effects of other kernels.

**Supplementary Table S1. Effects of Session and Run**

|  | **MID Anticipation** | | **SR Outcome** | |
| --- | --- | --- | --- | --- |
|  | VS Activation | BRS Correlation | VS Activation | BRS Correlation |
| Intercept | 0 (0.105) | 0 (0.107) | 0 (0.111) | 0 (0.105) |
| Session | -0.113 (0.108) | 0.124 (0.109) | -0.039 (0.115) | -0.156 (0.109) |
| Run | 0.080 (0.108) | 0.083 (0.105) | -0.025 (0.105) | 0.010 (0.108) |
| Mean Framewise Displacement | 0.047 (0.109) | -0.205 (0.107) | -0.142 (0.109) | 0.005 (0.109) |

**Supplementary Figure S3. Pre-Mood Induction Reward Response across Time**

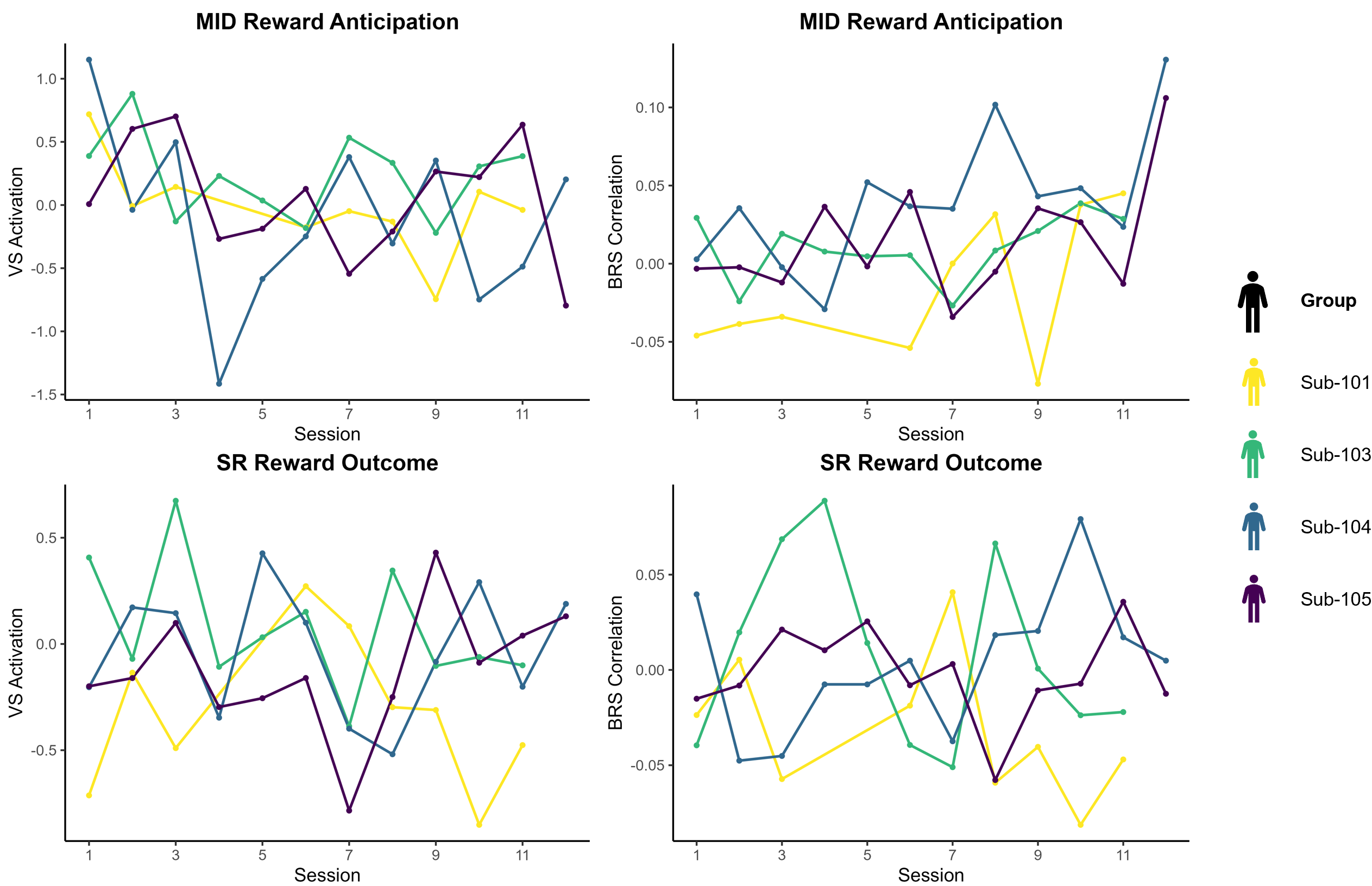

**Supplementary Figure S4. Intraindividual Variance Explained**

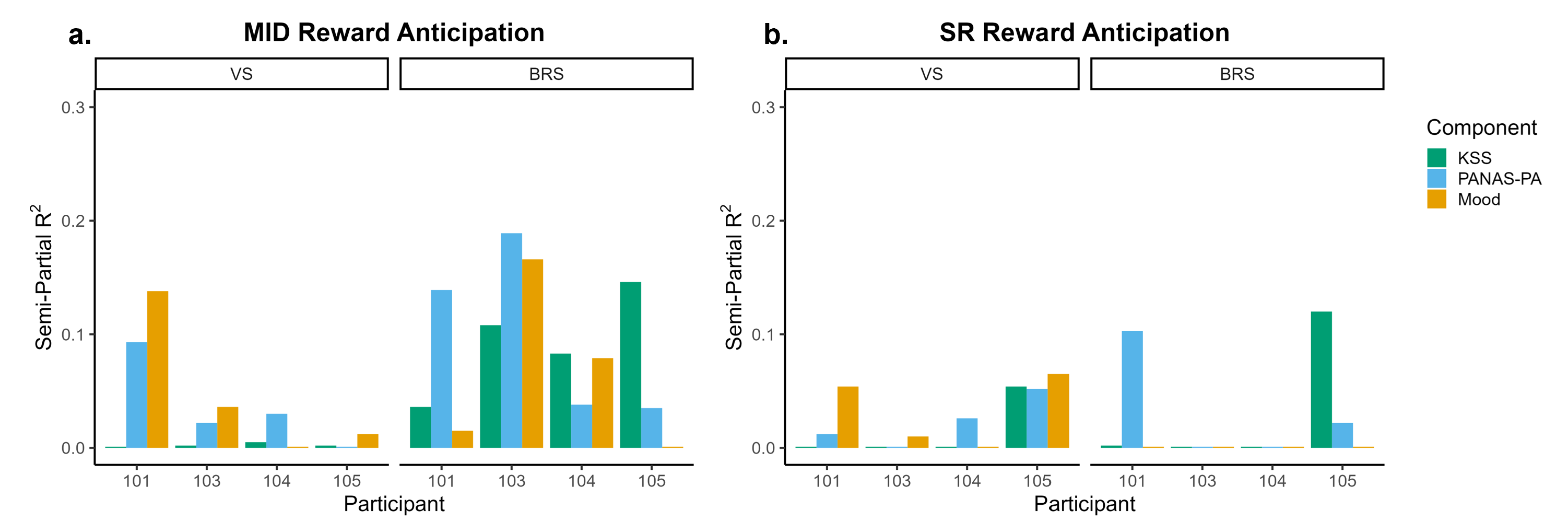
